# Rhythmicity Revisited: Evidence from Robust Behavioral Oscillations

**DOI:** 10.64898/2025.12.24.696378

**Authors:** Mengting Xu, Esperanza Badaya, Mehdi Senoussi, Tom Verguts

## Abstract

Rhythmic fluctuations in neuronal excitability, called neural oscillations, pervade brain activity. If these oscillations have a functional role in behavior, they should also be expressed in a behavioral signature. However, identifying such a behavioral signature has remained controversial: evidence for behavioral oscillations is not yet firmly established, and aperiodic components can be misidentified as rhythmic. To test their existence while controlling for aperiodic dynamics, we applied a state-of-the-art autoregressive (AR) approach on two datasets: one published dataset, and a new dataset collected with a revised dense-sampling design. In both datasets, AR modeling recovered reliable behavioral oscillations in the theta band. Reanalyses of the published dataset also reproduced the reported theta shift with task demands, with more difficult tasks showing slower theta. In contrast, despite clear rhythmicity, the new dataset showed no consistent frequency shift across conditions. Overall, this study establishes rigorous and robust detection of behavioral oscillations. It also suggests that the frequency modulation of behavioral oscillations by task difficulty requires further study.

## Introduction

A central challenge for cognition is minimizing interference between concurrently processed information. One influential proposal is that the brain accomplishes this by using neural oscillations, which function as an internal clock to segregate competing representations into separate temporal windows, thereby gating information flow and reducing cross-talk^1–4^. Nonetheless, some have questioned whether such oscillations are necessary or even genuinely present, arguing that cognitive operations (i.e., feature binding) can be supported by alternative, non-rhythmic dynamics such as event-like, aperiodic transients in spiking activity or more sustained changes in firing rates, without relying on oscillatory synchronization^5–7^, or that feature binding is not even necessary^8^. We address this debate by providing new evidence consistent with the oscillatory segregation account.

Electrophysiological studies have extensively documented how oscillatory dynamics are engaged and modulated by cognitive demands. Theta rhythms, for example, support multi-item working memory maintenance, slowing to accommodate more items^9–11^, whereas alpha rhythms often accelerate under higher cognitive load^12^. Such frequency shifts also contribute to the refined sampling of attentional^13,14^ and perceptual information^15,16^, which typically operate at distinct rhythms and adjust with task demands. This adaptability makes oscillatory frequency a promising marker of cognitive flexibility.

At a finer timescale, phase-dependent effects provide further support for this temporal account. The instantaneous phase of ongoing rhythms predicts performance in perception^17,18^, attention^13,19^, and memory^20,21^, reinforcing the view that neural computations are timed by oscillatory activity. A key question, however, follows: if this rhythmic organization is indeed fundamental to cognition, does its signature translate to overt, measurable behavior?

A growing body of research reports such periodic fluctuations in performance, known as behavioral oscillations, which provide compelling evidence for the role of neural rhythms in cognition. Indeed, if oscillations are a mere epiphenomenon of neural processing, it is unclear why or how they would have downstream effects on behavior. Studies using dense sampling design have revealed theta-rhythmic (∼4-8 Hz) oscillations in visual working memory accuracy^22^, while others have shown that response times systematically vary with the phase of ongoing theta and beta oscillations^23^. If behavioral oscillations are functionally meaningful, their properties (e.g., power or frequency) may also adapt to task requirements.

Initial evidence for a behaviorally relevant frequency adaptation came from Senoussi et al.^24^, who revealed that behavioral theta rhythms shift toward an optimal frequency as task demands change. For example, theta frequency slowed down when task rules became more difficult. However, that study relied on spectral methods with limited power to separate true periodicities from artifacts of aperiodic dynamics. Moreover, the authors did not explicitly test for oscillations in the first place, leaving open questions about the robustness of the reported effects. This methodological gap motivates our use of a participant-level autoregressive model^25^. Its key advantage lies in explicitly isolating the periodic component by characterizing and removing each participant’s aperiodic structure at the individual level, thereby minimizing the risk of misinterpreting noise as behavioral rhythms. Moreover, compared with a group-level autoregressive model^26^, this pipeline does not assume phase alignment across participants, which also helps preserve individual rhythmic signals.

Using this more rigorous approach, we examined two datasets. In Experiment 1, we reanalyzed the published dataset from Senoussi et al.^24^, which allowed us to test whether behavioral theta oscillations remain robust after controlling for aperiodic structure, and whether their frequency still varies with task difficulty. This reanalysis not only offers direct evidence for rhythmicity in behavior but also verifies the previously reported shifts in theta frequency. In Experiment 2, we aimed to replicate this result and used this occasion to address a limitation of the original paradigm. In that paradigm, the observed theta shift could reflect the influence of cue ambiguity or cue entropy (i.e., some cues contained two different symbols and other cues one), rather than being driven by task demand. Therefore, in Experiment 2, we designed a new paradigm where each cue showed exactly two different stimuli, but otherwise with the same cue meanings as in Experiment 1. Thus, we were able to dissociate cue ambiguity or entropy from task difficulty.

Together, these experiments pursue two main objectives: first, to provide stronger, methodologically robust evidence that behavior itself is rhythmic; and second, to test whether the frequency of behavioral oscillations is a general and reliable marker of cognitive demand. This is both a stringent test of the reliability of behavioral time courses and a deeper look at how the brain uses internal rhythms to organize behavior.

## Methods

### Experiment 1

#### Participants

Thirty-nine healthy adults (27 female; age 23.7 ± 4.5) took part in the study. All had normal or corrected-to-normal vision and no history of neurological or psychiatric disorders. Five participants were excluded based on the same data quality criteria as in the original study: two completed fewer than 5 blocks; one contributed fewer than 200 trials after eye-tracking-based trial rejection; one showed poor overall behavioral performance (<50% accuracy); and one was left-handed. The final sample comprised 34 participants, which provided at least 80% power to detect effect sizes ranging from moderate to large (Cohen’s f > 0.25, using repeated-measures ANOVA). The study was approved by the local ethics committee (Faculty of Psychology and Educational Sciences, Ghent University).

#### Experimental apparatus

Stimuli were presented using the PsychoPy toolbox (Python 2.7) on a 24-inch LCD monitor (1280 × 1080 pixels; 60 Hz) at a viewing distance of 60 cm, with head position stabilized by a combined chin- and forehead-rest. The testing room was dimly lit and kept at a constant illumination to minimize visual distractions and maintain stable ambient lighting during eye tracking.

#### Experimental stimuli and paradigm

The task required participants to discriminate the orientation of a cued target grating (Fig. 1A). On each cue display, two letters were presented simultaneously with a size of 0.75° visual angle (vertical eccentricity of 1° visual angle): one above the fixation cross indicated which of the forthcoming gratings would be the target, and one below the fixation cross indicated which hand to use for the response. Cue letters were drawn from ‘L’ and ‘R’, yielding four combinations (LL, RR, LR, RL). LL and RR were defined as the easy (congruent) rule condition, whereas LR and RL constituted the difficult (incongruent) rule condition. On the subsequent target display, two sinusoidal gratings were presented (size of 5° visual angle; 10% contrast; 3 cycles per degree) on a mid-grey background, centered at 5° horizontal eccentricity to the left and right of fixation. The tilt angle was individually titrated using a staircase procedure (see below for details) to equate perceptual difficulty and avoid ceiling accuracy. Participants were requested to report the tilt of one of two gratings as being clockwise (CW) or counter-clockwise (CCW) from the vertical axis, using the index or middle finger of one of both hands (Fig. 1C). On top of this, in order to assess rhythmic fluctuations in behavioral performance, we employed a dense-sampling design with evenly spaced inter-trial intervals (ISIs). ISIs ranged from 1700 to 2200ms in 50ms steps (11 levels) and were randomly assigned on each trial. Then, by aggregating accuracy and response time across ISIs (Fig. 1D), we were able to quantify oscillatory structure in behavior.

**Figure 1.**
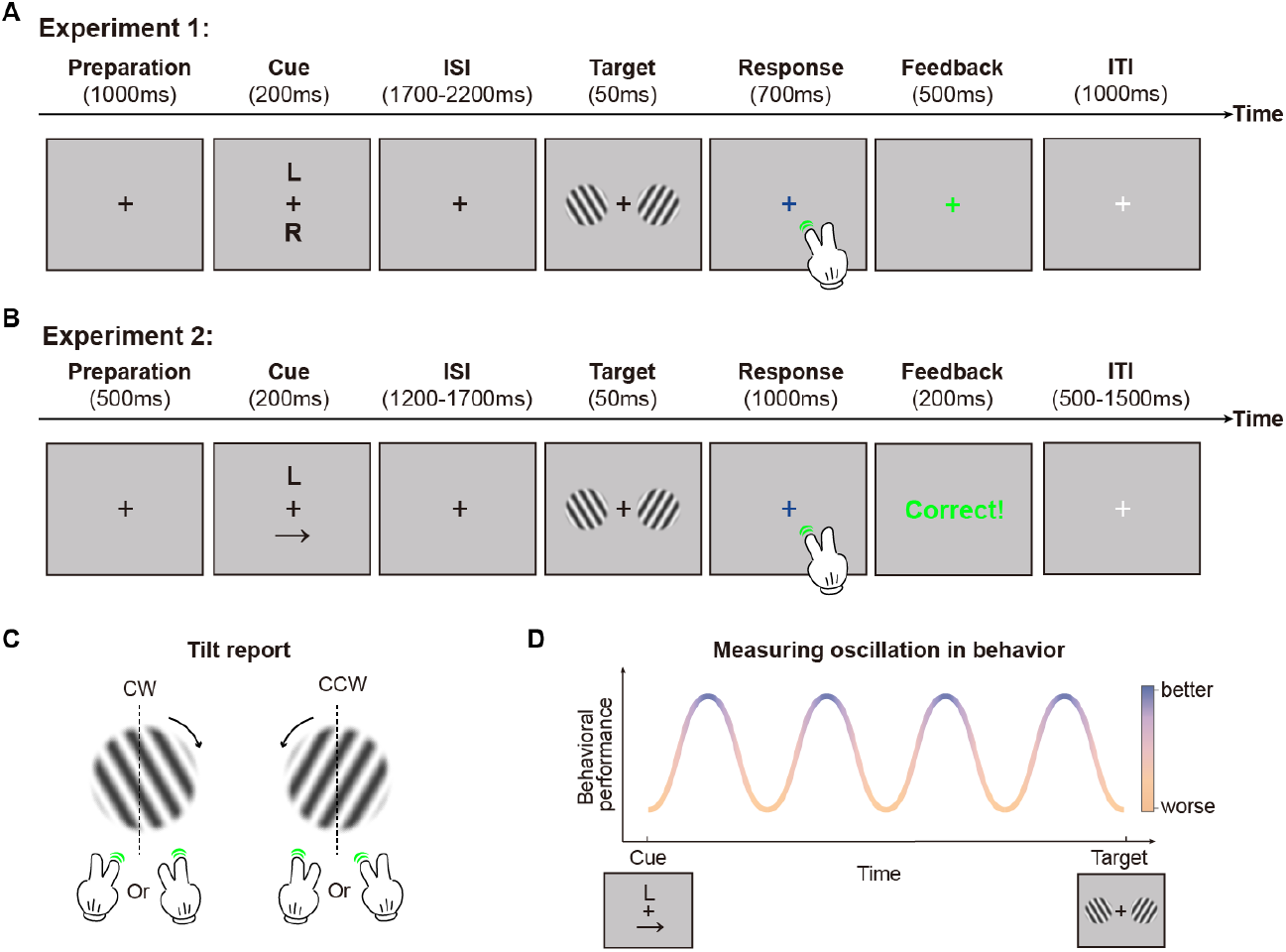
Paradigm. (A) Illustration of a single trial in Experiment 1. (B) Illustration of a single trial in Experiment 2. (C) Task display. Participants reported whether the target grating was tilted CW or CCW relative to vertical, using the index or middle finger of the cued hand. (D) Behavioral performance (accuracy/reaction time) was sampled across cue–target intervals, yielding a time course on which rhythmic fluctuations can be assessed.

#### Single-trial sequence

Each trial began with a preparatory interval (1000 ms), followed by the cue display (200ms). After a variable ISI, the target gratings appeared (50ms). During the response window, the fixation cross changed from black to blue to indicate that participants should respond (maximum response time: 700 ms). Feedback was then provided for 500 ms via the fixation cross (green for correct, red for incorrect). Trials without a response were marked as misses and re-presented at the end of the block; the task was paused at the same time, and a prompt asked whether a break was needed. If no break was required, participants pressed the ‘space’ to proceed. An inter-trial interval (ITI) of 1000 ms then preceded the next trial.

#### Experimental procedure

Each participant first completed a practice block (80 trials; details see Senoussi et al.^24^), followed by a staircase procedure (80 trials) to estimate an individual tilt level for use in the main task. Stimulus settings and timings were identical to the main task, with only grating orientation varied on a trial-to-trial basis, and trials were sampled across all cue types and ISIs to avoid ceiling performance. We used a one-up, two-down procedure with the tilt level initialized at 7° with 3° steps (ranging from 0.5 to 30°; minimum step 0.1°). The step size was halved on every alternate reversal after the second reversal. Reversals happened at transitions from a run of correct responses to an error, or vice versa: after an error following correct responses, tilt increased (became easier); after a correct response following errors, tilt decreased (became harder). The tilt values carried into the main task were the mean of the final ten staircase values. The main task comprised 5–8 blocks, depending on participant pace (those with fewer missed trials completed more blocks).

#### Data preprocessing and cleaning

The published behavioral dataset was acquired with an eye-tracker (acquisition and processing details are provided in Senoussi et al.^24^). In the current pipeline, we likewise rejected trials based on gaze position (epoched from cue onset to target onset) relative to the fixation cross, using a 1.5° threshold as in the original study. After data cleaning, trials were binned by condition (cue type) and by ISI. We then computed mean accuracy and mean RT within each bin, generating condition-specific time courses of behavioral performance across ISIs, that is, behavioral oscillations.

#### Spectral analysis of behavioral oscillations

We used a state of the art autoregressive approach^25^ to estimate the spectral representation of behavioral oscillations. This method starts by fitting a first-order autoregressive (AR(1)) model to each participant’s ISI-binned time course^25^:

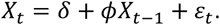

We estimated the parameters (constant *δ*, autoregressive coefficient *ϕ*) per subject and simulated 1000 surrogate series of length *T* (with Gaussian noise *ε*_*t*_) for each subject. These surrogates capture non-rhythmic trial-by-trial dependencies (e.g., priming). We then computed fast Fourier transform (FFT) amplitude spectra for the empirical series and all surrogates after zero-padding each time series with 32 samples (adding 16 zeros before and 16 after). For each frequency *f*, we defined the participant-level deviation spectrum as:

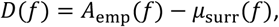

where *A*_*emp*_ is the empirical amplitude and *μ*_surr_(*f*) is the mean amplitude across 1000 surrogates. Positive *D*(*f*) indicate the presence of oscillatory activity.

To assess whether a common behavioral rhythm was present at the group level, we evaluated each frequency *f* with a one-tailed one-sample *t*-test across participants (mean *D*(*f*) > 0). *p*-values were then adjusted across frequencies using Benjamini–Hochberg FDR correction (*q* = 0.05). We report FDR-adjusted *p*-values and highlight frequencies that survive FDR as group-level rhythmic peaks.

### Experiment 2

#### Participants

Forty-two adults (32 females; mean age = 25.83 ± 3.25) were recruited. All had normal or corrected-to-normal vision and no history of neurological or psychiatric disorders. Two participants were excluded (one for low accuracy and one for reversed response mapping), and fourteen were excluded based on predefined eye-movement criteria (see Data preprocessing and cleaning). The final sample comprised twenty-six participants, providing at least 80% power to detect experimental effect sizes ranging from moderate to large (Cohen’s f > 0.25, using repeated-measures ANOVA). The study was approved by the local ethics committee (Faculty of Psychology and Educational Sciences, Ghent University).

#### Experimental apparatus

Stimuli were presented with the PsychoPy toolbox (Python 3.10) on a 24-inch LCD monitor (1920 × 1080 pixels; 60 Hz) at a viewing distance of 93 cm, with head position stabilized by a combined chin and forehead rest. The testing room was dimly lit and kept at a constant illumination (∼30 lux, measured with an illuminance meter) to minimize visual distractions and maintain stable ambient lighting during eye tracking. To ensure that task difficulty was not influenced by gaze position, participants were instructed to maintain fixation on a central fixation cross (size 0.30 °). Eye movements of the dominant eye (monocular tracking) were recorded continuously with an EyeLink 1000 Plus desktop-mounted system (SR Research) at 1000 Hz, and all eye-tracking procedures were controlled via PsychoPy scripts. A 13-point calibration and validation procedure was performed at the beginning of each block, followed by a drift correction once per block (every 25 trials).

#### Experimental stimuli and paradigm

To ensure that any frequency shifts in behavioral oscillations reflected task difficulty rather than stimulus semantics, we modified the original design (see Experiment 1) to use letter–arrow cues instead of letters alone. Cue symbols were drawn from L, R, ←, →, yielding 16 two-symbol combinations. On each cue display, the instructions matched the original paradigm, with the upper symbol indicating the target location (left or right) and the lower symbol indicating the response hand. Cues were grouped by spatial congruency into three conditions: SS (same symbols: LL, RR, ←←, →→), SSiDS (same side, different symbols: L←, ←L, R→, →R), and DSi (different sides: L→, LR, R←, RL, ←R, ←→, →L, →←). To equate exposure across conditions, each participant viewed a counterbalanced subset of four DSi pairs (Fig. 1B). The two cue symbols were presented to the left and right of fixation at ±0.75° horizontal eccentricity, with a size of 0.75°. On the target display, a pair of sine-wave Gabor gratings was presented symmetrically at ±7.5° horizontal eccentricity (size 5°, contrast 0.2, spatial frequency 2 cycles per degree) on the mid-grey background. The task was identical to Experiment 1: participants reported whether the target grating was tilted clockwise (CW) or counterclockwise (CCW) relative to vertical, using the index or middle finger of the instructed hand (Fig. 1C).

#### Single-trial sequence

Each trial began with a 500 ms preparation baseline, followed by presentation of the two cue symbols for 200 ms. A cue–target interval of 1200–1700 ms (step size 50 ms) then ensued. The target display was presented for 50 ms, and participants subsequently had 1000 ms to report whether the target grating was tilted clockwise or counterclockwise. Accuracy feedback was shown for 200 ms (green ‘Correct!’ for correct responses; red ‘Incorrect!’ for incorrect responses), after which an ITI of 500–1500 ms (step size 250 ms) preceded the next trial. Trials with no response were marked as misses and were re-presented at the end of the current block.

#### Experimental procedure

The experiment comprised two sessions on separate days. In each session, participants began with a brief ROI familiarization phase (∼60 s) with gaze-contingent auditory feedback to encourage stable central fixation. During this phase, a central fixation marker (green dot; 20×20 pixels) was presented together with a circular ROI (size 1.5°), and a beep was triggered whenever the gaze deviated outside the ROI. In Session 1, participants then completed a practice block (100 trials) to familiarize themselves with the task (target duration 100 ms; response window 1500 ms), followed by a staircase block (100 trials) identical to that in Experiment 1. The resulting individual tilt angle from the staircase procedure was then used in the main task for both sessions. In Session 2, participants completed a shorter practice block (25 trials) and then proceeded directly to the main task. The main task across the two sessions comprised 1056 trials in total (428 in Session 1 and 628 in Session 2) and was organized into blocks of 25 trials each.

#### Data preprocessing and cleaning

We assessed fixation quality on every trial during the cue-to-target interval. A trial was marked as poor quality if gaze deviated by more than ∼2° from the central fixation cross at any time during this interval. Participants were excluded when more than 50% of their trials failed this criterion across the two sessions; in total, 14 participants were removed for low eye-tracking quality. At the session level, we required accuracy to be reliably above chance using a one-sided binomial test against 0.5 (α = 0.05). Two sessions (from two participants) did not pass this check and were excluded. The final dataset comprised all remaining sessions and included only valid trials (all miss trials excluded). Valid trials were then binned by condition (SS, SSiDS, and DSi) and ISI, and we computed mean accuracy and mean reaction time within each bin to prepare for the estimation of behavioral oscillations.

### Spectral analysis of behavioral oscillations

Identical to Experiment 1.

## Results

### Experiment 1

#### Replicating theta-rhythmic modulation of behavior

All subsequent analyses were conducted on a robust dataset comprising 34 participants, following the exclusion of five individuals based on pre-defined data quality criteria (see Methods). Crucially, the oscillatory effects reported below were robust to these participant removal criteria. We began by verifying rhythmicity in the behavioral time courses of this published dataset by quantifying deviation spectra from participant-specific surrogate spectra (see Methods). In accuracy, clear oscillatory structure was evident across all four conditions (RR, LL, LR, RL; Fig. 2A, top), with somewhat stronger evidence under the difficult rules (LR and RL). Specifically, in the two easy rules (RR and LL) a single frequency bin reached *t*-test significance but did not pass FDR correction (3.75Hz in RR and 6.25Hz in LL). By contrast, in LR five bins were statistically significant (3.75Hz, 4.38Hz, 6.25 Hz, 6.88 Hz, and 7.50 Hz), of which four remained significant after FDR correction (*p*_*FDR*_ < .01 at 3.75Hz, 4.38Hz and 6.25Hz; *p*_*FDR*_ = .01 at 6.88Hz); similarly in RL, *t*-tests were significant in four bins, with three retaining significance after FDR correction (*p*_*FDR*_ < .01 at 3.75Hz, 4.38Hz and 5Hz). For the RT, rhythmicity was likewise observed in all conditions (Fig. 2A, bottom): in RR, the *t*-test was significant in two bins (4.38Hz and 3.75Hz), of which one bin was FDR-corrected significant (*p*_*FDR*_ = .01 at 3.75Hz); in LL, one bin reached uncorrected significance (5Hz); in LR, three bins were significant (3.75Hz, 4.38Hz and 5 Hz), of which two remained significant after FDR correction (*p*_*FDR*_ < .001 at 3.75Hz ; *p*_*FDR*_ < .01 at 4.38Hz); and in RL, one bin met the FDR-corrected threshold (*p*_*FDR*_ = .01 at 3.75Hz). Taken together, behavioral performance exhibits theta-band rhythmic fluctuation, which supports a true oscillatory process, possibly on top of aperiodic structure.

**Figure 2.**
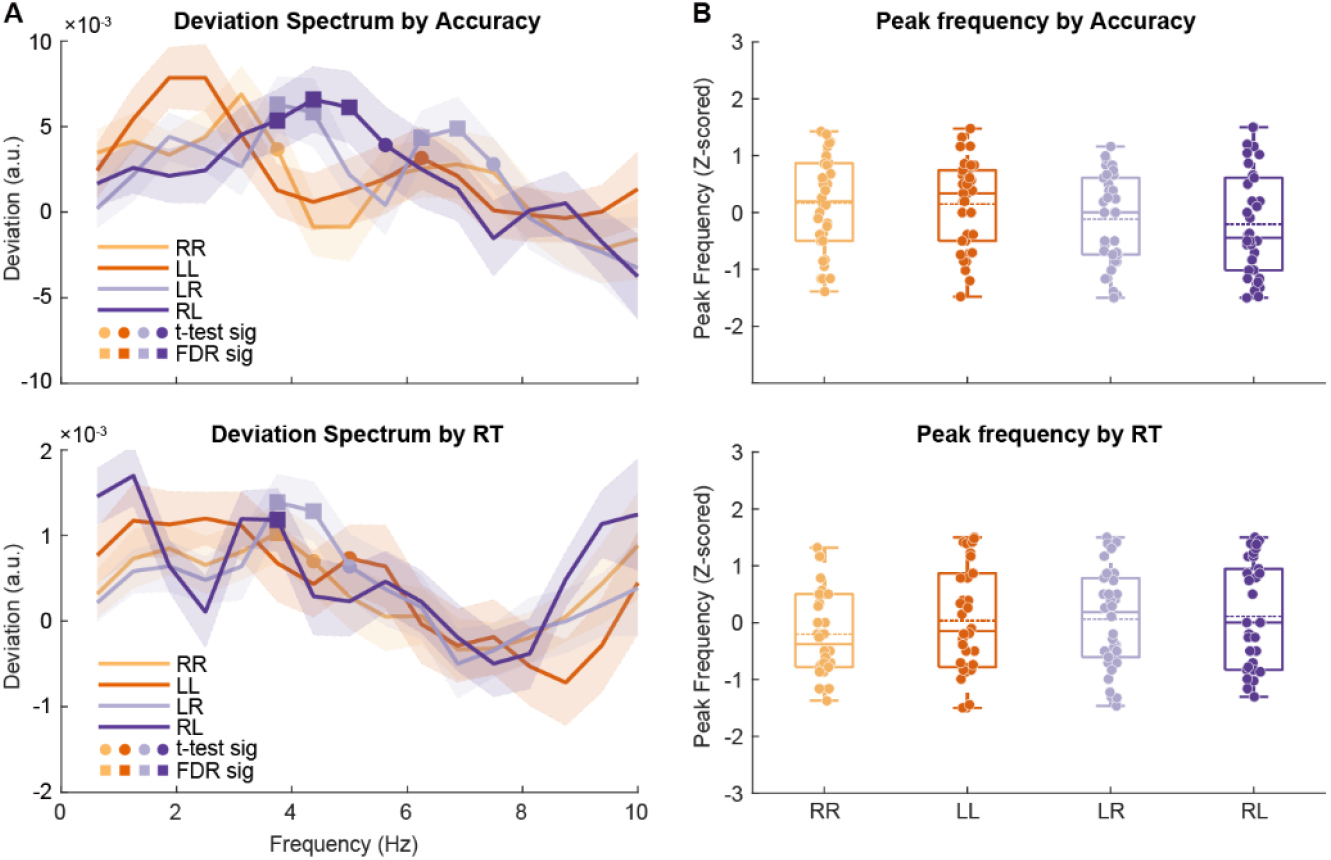
Results of Experiment 1. (A) Group-mean deviation spectra for accuracy (top) and reaction time (bottom). Curves correspond to the four conditions (RR, LL, LR, RL); shaded areas indicate ± s.e.m. Filled circles mark frequencies significant by one-tailed one-sample t-tests across participants (*p* < 0.05); filled squares mark frequencies surviving Benjamini– Hochberg FDR correction across frequencies (*q* = 0.05). Statistical analysis was restricted to the theta-frequency range (3.75 - 7.50 Hz). (B) Distributions of individual theta peak frequencies (z-scored across all conditions) for accuracy (top) and reaction time (bottom). Box plots show the median and interquartile range (whiskers, 1.5× IQR); points indicate individual participants; solid horizontal lines indicate the mean peak frequency.

#### Reproducing the difficulty-dependent shift in peak frequency

A key observation in the earlier publication^24^ is the theta-frequency shift according to task demands, which we sought to replicate under the AR-model framework. Accordingly, we defined the peak frequency for each participant and condition as the frequency bin with maximal deviation and compared the four conditions using a 2 (hands) × 2 (target locations) linear mixed-effects model (LMM) with participants as a random intercept, applied separately to accuracy and RT. Consistent with the original report, in accuracy, we observed a reliable difficulty-dependent shift in peak frequency: peak theta frequency significantly decreased from the easy rules (RR and LL) to the difficult rules (LR and RL; *β* = −.16, *p* = .03; Fig. 2B, top). Parallel analyses using RT-based peaks found no reliable modulation by rule difficulty (*β* = .09, *p* = .25; Fig. 2B, bottom). Taken together, these results indicate that the frequency shift remains robust under the AR framework, which further supports that rule-dependent slowing of theta tracks cognitive-control demands at the behavioral level.

### Experiment 2

#### Behavioral performance

The final sample included 26 participants following predefined eye-tracking and binomial screening (see Methods). Of these, 24 participants contributed 1056 trials across both sessions, while 2 participants contributed 628 trials from Session 2 only. Critically, all reported behavioral effects remained robust to variations in participant exclusion criteria. Accuracy followed the expected prediction, SSiDS ≈ SS > DSi (means: SS = 79.16%, SSiDS = 79.94%, DSi = 70.69%; Fig. 3A). A repeated-measures ANOVA confirmed a condition effect, *F*_(2,50)_ = 15.88, *p* < .001, *η*_*p*_^*2*^ = .39. Paired *t* tests (two-tailed) showed lower accuracy for DSi relative to SS (*t*_*(25)*_ = 3.85, *p* < .001, *d*_*z*_ = 0.76) and SSiDS (*t*_*(25)*_ = 4.27, *p* < .001, *d*_*z*_ = 0.84), whereas SS and SSiDS did not differ (*t*_*(25)*_ = −1.23, *p* = .23, *d*_*z*_ = −0.24). RT showed a similar pattern, SSiDS > SS > DSi (means: SSiDS = 565.37ms, SS = 572.24ms, DSi = 590.83ms; Fig. 3B). A repeated-measures ANOVA also showed a condition main effect, *F*_*(2,50)*_ = 18.78, *p* < .001, *η*_*p*_^*2*^ = .43, with SSiDS faster than SS (*t*_*(25)*_ = 2.88, *p* < .01, *d*_*z*_ = 0.56) and both faster than DSi (SS vs DSi: *t*_*(25)*_ = −3.37, *p* < .01, *d*_*z*_ = −0.66; SSiDS vs DSi: *t*_(25)_ = −5.79, *p* < .001, *d*_*z*_ = −1.14). Together, the behavioral patterns show that DSi cues place the greatest control demands.

**Figure 3.**
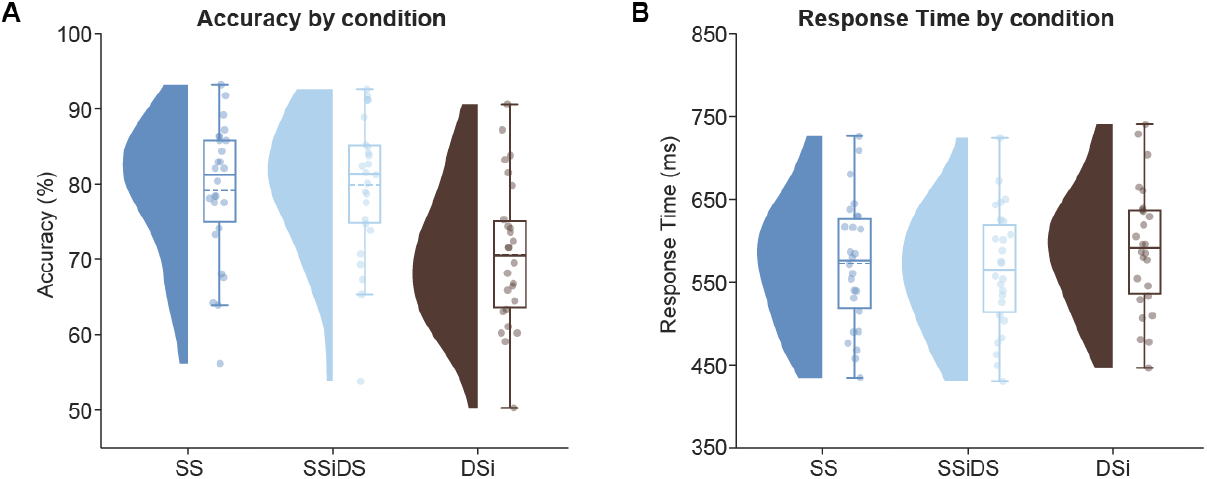
Group-level behavioral performance in Experiment 2. (A) Accuracy across conditions. Violin plots show the across-participant distribution; box plots indicate the median and interquartile range (whiskers, 1.5× IQR); points indicate individual participants; solid horizontal lines indicate the mean values. (B) Similar to (A), but for Response time.

#### Replicating theta-rhythmic modulation of behavior

As in Experiment 1, we analyzed accuracy and RT by densely sampled cue–target intervals and, for each condition, computed deviation spectra relative to participant-level autoregressive surrogates to control aperiodic structure^25^. Accuracy spectra exhibited theta-band rhythmic fluctuation (∼4–8 Hz; Fig. 4A, top). In the SS condition, three theta peaks were observed: a peak at 5.63 Hz that survived FDR correction (*p*_*FDR*_ = .03), and two peaks at 5 Hz and 6.25 Hz that were significant in *t*-tests but did not survive FDR correction. In the DSi condition, two peaks reached *t*-test significance (3.75 Hz and 4.38 Hz), but neither survived FDR correction. No peak was significant in the SSiDS condition. RT spectra showed a similar pattern (Fig. 4A, bottom): Two frequencies were significant after FDR-correction in SS (*p*_*FDR*_ = .04 at 5 Hz and 5.63 Hz); two theta frequencies reached *t*-test significance in SSiDS (5.63 Hz and 6.25 Hz) but did not survive FDR-correction. Together, these analyses reveal a theta-band component in behavior, consistent with a common oscillatory mechanism across SS conditions.

**Figure 4.**
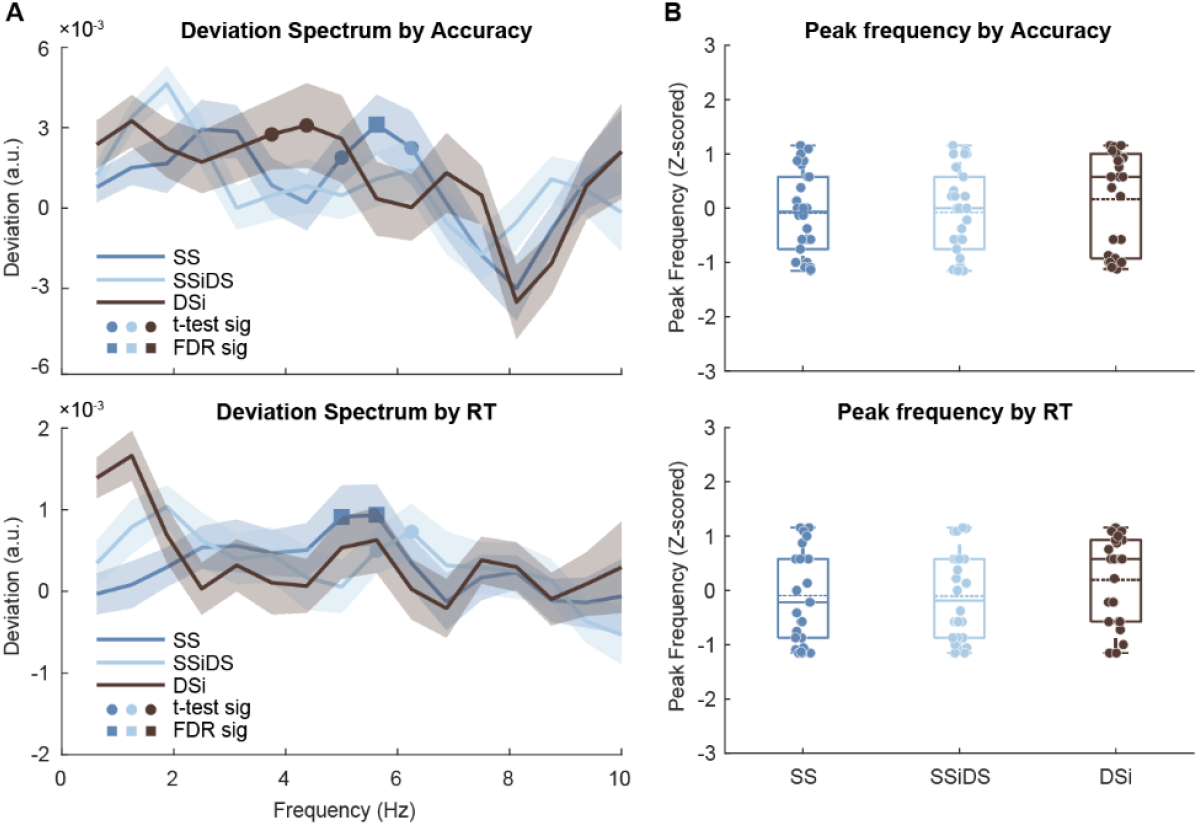
Results of Experiment 2. (A) Group-mean deviation spectra for accuracy (top) and reaction time (bottom). Curves correspond to the three conditions (SS, SSiDS, and DiS); shaded areas indicate ± s.e.m. Filled circles mark frequencies significant by one-tailed one-sample t-tests across participants (*p* < 0.05); filled squares mark frequencies surviving Benjamini– Hochberg FDR correction across frequencies (*q* = 0.05). Statistical analysis was restricted to the theta-frequency range (3.75 - 7.50 Hz). (B) Distributions of individual theta peak frequencies (z-scored across all conditions) for accuracy (top) and reaction time (bottom). Box plots show the median and interquartile range (whiskers, 1.5× IQR); points indicate individual participants; solid horizontal lines indicate the mean peak frequency.

#### No difficulty-dependent shift in peak frequency

In order to assess whether the reported theta-frequency shift also generalizes in this new dataset, we applied the same analysis pipeline as Experiment 1 to extract, for each participant and condition, the theta peak defined as the frequency bin with maximal deviation in the accuracy and RT spectra. A linear mixed-effects model testing the main effect of condition revealed no differences in peak frequency across SS, SSiDS, and DSi for either accuracy (Type-III ANOVA with Satterthwaite df: *F*_*(2,78)*_ = 0.79, *p* = .46; Fig. 4B, top) or RT (*F*_*(2,78*)_ = 1.14, *p* = .32; Fig. 4B, bottom). We then examined whether peak frequency predicts performance using generalized linear mixed models with accuracy and RT as dependent variables to replicate another analysis from Senoussi et al^24^. Peak frequency did not predict accuracy (*β* = -.08, *p* = .19), nor RT (*β* = .006, *p* = .59). Collectively, this new dataset provides no evidence that task difficulty shifts the behavioral theta peak.

## Discussions

The present study revisited accounts of rhythmic cognition by using a state-of-the-art autoregressive (AR) surrogate approach^25^ on two behavioral datasets. In Experiment 1 (Fig. 1A), a reanalysis of a previously published dataset revealed robust theta-band oscillations in both accuracy and response time (Fig. 2A), even after explicitly modelling and removing each participant’s aperiodic temporal structure. Using this more conservative pipeline, we replicated the earlier finding^24^ that the peak theta frequency of accuracy slows under more demanding rule conditions (Fig. 2B, top). In Experiment 2, we used a modified cue design that separated cue ambiguity from task difficulty (Fig. 1B) and again observed theta-band rhythmicity in behavioral performance (Fig. 4A). In contrast to Experiment 1, however, we did not find reliable frequency shifts as task demands changed (Fig. 4B). Taken together, these results strengthen the evidence that behavioral time courses contain genuine theta-band oscillations, but find no additional evidence that frequency of behavioral rhythms shifts with task difficulty.

Frequency shifts have been repeatedly observed in EEG^15,16,27–31^, but the evidence for behavioral shifts remains limited. The discrepancy in frequency-shift findings between experiments calls for a careful consideration of limitations. In both experiments, we employed a dense sampling design. This design is powerful for detecting rhythmicity^22,32,33^, but it provides only a limited number of data points per participant within each condition and each interval. This is particularly problematic for detecting subtle shifts in peak frequency, which are likely to be small and to vary across individuals^12,34^. Thus, the null result in Experiment 2 does not necessarily imply that frequency modulation is absent; it may also reflect insufficient within-participant data to estimate such shifts with confidence. In addition, the AR(1) surrogate model is itself a relatively simplified assumption about behavioral temporal dynamics^25,26^. Real behavioral time courses may contain higher-order or nonlinear dependencies that are not captured by this model, which could in turn bias the surrogate-derived baseline^35^ and thereby affect estimates of true periodicity.

These limitations point to clear directions for methodological improvement. One straightforward step is to adopt multi-session designs (see Hachenberger et al.^36^), in which participants complete many more trials spread across multiple sessions. Such designs (with more data points per subject) would allow more precise estimation of individual oscillatory profiles (e.g., peak frequency; see Drewes et al.^37^) and could also reduce the impact of between-subject variability. In parallel, efforts that combine data from multiple laboratories or multiple datasets through meta-analysis^38^ could help clarify whether frequency shifts reliably appear under specific task demands^16,39^, sampling schemes^40^, or analysis pipelines (to separate periodic and aperiodic components)^25,26,41^. A meta-analytic approach would also allow formal quantification of effect sizes and heterogeneity across paradigms, which is difficult to assess from single studies^42,43^.

Moreover, these results raise a broader mechanistic question: how can oscillations help keep concurrent streams of information apart during cognition? Many theories converge on the idea that a core function of oscillations is to minimize interference between representations (e.g., ‘Communication through Coherence’)^1,44^. In such accounts, oscillatory dynamics organize information processing into recurring temporal windows, with distinct phases within a single cycle assigned to different representations^21,45,46^. This temporal segregation helps prevent overlap between competing representations^47,48^, while still supporting effective feature binding^49,50^. In our paradigm, participants must switch between paying attention to the left and ride side information, and responding with the left and right hands. Because all information will be partially activated across trials, temporal binding of the currently-relevant task processes (e.g., for cue LR, left stimulus and right hand) via rhythmic alternation can help reduce representational interference and help to address the binding problem^51,52^. Our observation of behavioral theta-frequency fluctuations in Experiment 1, together with the persistence of rhythmicity in Experiment 2 even without a reliable frequency shift, supports the notion that such phase-based segregation is a fundamental strategy for minimizing interference in cognition.

In summary, applying rigorous and robust (AR-based) state-of-the-art analyses to both a published and a new dense-sampling dataset provides converging evidence that cognition is intrinsically rhythmic; although adaptive modulation of such rhythms remains to be investigated using different tasks or difficulty manipulations and more powerful statistical and analytical approaches. These findings suggest that the brain organizes behavior not as a continuous flow, but as a series of discrete, rhythmically gated operations that support efficient processing by minimizing interference in cognition.

## Data availability

All data and research materials for Experiment 1 are available at https://osf.io/nwh87/?view_only=b11ee1f860804da582c816fe8acdecad.

All data and research materials for Experiment 2 are available at https://osf.io/3k2r5/?view_only=949efc2399f04e079d8964037243692d.

## Code availability

All code for Experiments 1 and 2 is available at https://github.com/CogComNeuroSci/Mengting_public/tree/main/rhythmic%20cognition.

## Acknowledgments

This work was supported by the China Scholarship Council (CSC; 202406750012) and the Fonds Wetenschappelijk Onderzoek - Vlaanderen (FWO; 1117026N). The funders had no role in the study design, data collection, analysis, or interpretation, the decision to publish, or the preparation of the manuscript.

## Author contributions

M.X., and M.S., designed the research, collected the data, performed data analysis, and wrote the manuscript. T.V. designed the research, performed data analysis, and wrote the manuscript. E.B. collected the data and wrote the manuscript. All authors discussed the results and commented on the manuscript.

## References

1. Fries, P. Rhythms for Cognition: Communication through Coherence. Neuron 88, 220–235 (2015).

2. Fries, P. Rhythmic attentional scanning. Neuron 111, 954–970 (2023).

3. Landau, A. N. & Fries, P. Attention Samples Stimuli Rhythmically. Current Biology 22, 1000– 1004 (2012).

4. VanRullen, R. Perceptual Cycles. Trends in Cognitive Sciences 20, 723–735 (2016).

5. Ray, S. & Maunsell, J. H. R. Do gamma oscillations play a role in cerebral cortex? Trends in Cognitive Sciences 19, 78–85 (2015).

6. Roelfsema, P. R. Solving the binding problem: Assemblies form when neurons enhance their firing rate—they don’t need to oscillate or synchronize. Neuron 111, 1003–1019 (2023).

7. Vinck, M. et al. Principles of large-scale neural interactions. Neuron 111, 987–1002 (2023).

8. Scholte, H. S. & De Haan, E. H. F. Beyond binding: specialization without segregation. Trends in Cognitive Sciences (2025).

9. Axmacher, N. et al. Cross-frequency coupling supports multi-item working memory in the human hippocampus. Proc. Natl. Acad. Sci. U.S.A. 107, 3228–3233 (2010).

10. Ratcliffe, O., Shapiro, K. & Staresina, B. P. Fronto-medial theta coordinates posterior maintenance of working memory content. Current Biology 32, 2121-2129.e3 (2022).

11. Wolinski, N., Cooper, N. R., Sauseng, P. & Romei, V. The speed of parietal theta frequency drives visuospatial working memory capacity. PLoS Biol 16, e2005348 (2018).

12. Haegens, S., Cousijn, H., Wallis, G., Harrison, P. J. & Nobre, A. C. Inter- and intra-individual variability in alpha peak frequency. NeuroImage 92, 46–55 (2014).

13. Fiebelkorn, I. C., Saalmann, Y. B. & Kastner, S. Rhythmic Sampling within and between Objects despite Sustained Attention at a Cued Location. Current Biology 23, 2553–2558 (2013).

14. Re, D., Tosato, T., Fries, P. & Landau, A. N. Perplexity about periodicity repeats perpetually: A response to Brookshire. Preprint at 10.1101/2022.09.26.509017 (2022).

15. Samaha, J. & Postle, B. R. The Speed of Alpha-Band Oscillations Predicts the Temporal Resolution of Visual Perception. Current Biology 25, 2985–2990 (2015).

16. Wutz, A., Melcher, D. & Samaha, J. Frequency modulation of neural oscillations according to visual task demands. Proc. Natl. Acad. Sci. U.S.A. 115, 1346–1351 (2018).

17. Mathewson, K. E., Gratton, G., Fabiani, M., Beck, D. M. & Ro, T. To See or Not to See: Prestimulus α Phase Predicts Visual Awareness. J. Neurosci. 29, 2725–2732 (2009).

18. Xu, M., Han, B., Chen, Q. & Shen, L. Auditory stimuli extend the temporal window of visual integration by modulating alpha-band oscillations. eLife 14, (2025).

19. Lakatos, P., Karmos, G., Mehta, A. D., Ulbert, I. & Schroeder, C. E. Entrainment of Neuronal Oscillations as a Mechanism of Attentional Selection. Science 320, 110–113 (2008).

20. Rizzuto, D. S., Madsen, J. R., Bromfield, E. B., Schulze-Bonhage, A. & Kahana, M. J. Human neocortical oscillations exhibit theta phase differences between encoding and retrieval. NeuroImage 31, 1352–1358 (2006).

21. Kerrén, C., Van Bree, S., Griffiths, B. J. & Wimber, M. Phase separation of competing memories along the human hippocampal theta rhythm. eLife 11, e80633 (2022).

22. Pomper, U. & Ansorge, U. Theta-Rhythmic Oscillation of Working Memory Performance. Psychol Sci 32, 1801–1810 (2021).

23. Abdalaziz, M., Redding, Z. V. & Fiebelkorn, I. C. Rhythmic temporal coordination of neural activity prevents representational conflict during working memory. Current Biology 33, 1855-1863.e3 (2023).

24. Senoussi, M. et al. Theta oscillations shift towards optimal frequency for cognitive control. Nat Hum Behav 6, 1000–1013 (2022).

25. Harris, A. M. & Beale, H. A. Detecting behavioural oscillations with increased sensitivity: A modification of Brookshire’s (2022) AR-surrogate method. eLife 14, (2025).

26. Brookshire, G. Putative rhythms in attentional switching can be explained by aperiodic temporal structure. Nat Hum Behav 6, 1280–1291 (2022).

27. Mierau, A., Klimesch, W. & Lefebvre, J. State-dependent alpha peak frequency shifts: Experimental evidence, potential mechanisms and functional implications. Neuroscience 360, 146–154 (2017).

28. Rodriguez-Larios, J. & Alaerts, K. Tracking Transient Changes in the Neural Frequency Architecture: Harmonic Relationships between Theta and Alpha Peaks Facilitate Cognitive Performance. J. Neurosci. 39, 6291–6298 (2019).

29. Riddle, J., Vogelsang, D. A., Hwang, K., Cellier, D. & D’Esposito, M. Distinct Oscillatory Dynamics Underlie Different Components of Hierarchical Cognitive Control. J. Neurosci. 40, 4945–4953 (2020).

30. Merholz, G., Grabot, L., VanRullen, R. & Dugué, L. Periodic attention operates faster during more complex visual search. Sci Rep 12, 6688 (2022).

31. Rassi, E. et al. Beta-band frequency shifts signal decisions in human prefrontal cortex. iScience 28, 113806 (2025).

32. Drewes, J., Zhu, W., Wutz, A. & Melcher, D. Dense sampling reveals behavioral oscillations in rapid visual categorization. Sci Rep 5, 16290 (2015).

33. Chota, S., Leto, C., Van Zantwijk, L. & Van Der Stigchel, S. Attention rhythmically samples multi-feature objects in working memory. Sci Rep 12, 14703 (2022).

34. Espenhahn, S., De Berker, A. O., Van Wijk, B. C. M., Rossiter, H. E. & Ward, N. S. Movement-related beta oscillations show high intra-individual reliability. NeuroImage 147, 175–185 (2017).

35. Schreiber, T. & Schmitz, A. Surrogate time series. Physica D: Nonlinear Phenomena 142, 346– 382 (2000).

36. Hachenberger, J. et al. Within-subject reliability, occasion specificity, and validity of fluctuations of the Stroop and go/no-go tasks in ecological momentary assessment. Behav Res 57, 29 (2024).

37. Drewes, J., Muschter, E., Zhu, W. & Melcher, D. Individual resting-state alpha peak frequency and within-trial changes in alpha peak frequency both predict visual dual-pulse segregation performance. Cerebral Cortex 32, 5455–5466 (2022).

38. Button, K. S. et al. Power failure: why small sample size undermines the reliability of neuroscience. Nat Rev Neurosci 14, 365–376 (2013).

39. Babu Henry Samuel, I., Wang, C., Hu, Z. & Ding, M. The frequency of alpha oscillations: Task-dependent modulation and its functional significance. NeuroImage 183, 897–906 (2018).

40. Kienitz, R., Schmid, M. C. & Dugué, L. Rhythmic sampling revisited: Experimental paradigms and neural mechanisms. Eur J of Neuroscience 55, 3010–3024 (2022).

41. Donoghue, T. et al. Parameterizing neural power spectra into periodic and aperiodic components. Nat Neurosci 23, 1655–1665 (2020).

42. Borenstein, M., Hedges, L. V., Higgins, J. P. T. & Rothstein, H. R. Introduction to Meta-Analysis. (Wiley, 2009). doi:10.1002/9780470743386.

43. Samaha, J. & Romei, V. Alpha-Band Frequency and Temporal Windows in Perception: A Review and Living Meta-analysis of 27 Experiments (and Counting). Journal of Cognitive Neuroscience 36, 640–654 (2024).

44. Fries, P. A mechanism for cognitive dynamics: neuronal communication through neuronal coherence. Trends in Cognitive Sciences 9, 474–480 (2005).

45. Siegel, M., Warden, M. R. & Miller, E. K. Phase-dependent neuronal coding of objects in short-term memory. Proc. Natl. Acad. Sci. U.S.A. 106, 21341–21346 (2009).

46. Lisman, J. E. & Jensen, O. The Theta-Gamma Neural Code. Neuron 77, 1002–1016 (2013).

47. Sauseng, P., Griesmayr, B., Freunberger, R. & Klimesch, W. Control mechanisms in working memory: A possible function of EEG theta oscillations. Neuroscience & Biobehavioral Reviews 34, 1015–1022 (2010).

48. Staudigl, T., Hanslmayr, S. & Bäuml, K.-H. T. Theta Oscillations Reflect the Dynamics of Interference in Episodic Memory Retrieval. J. Neurosci. 30, 11356–11362 (2010).

49. Gray, C. M. The temporal correlation hypothesis of visual feature integration: still alive and well. Neuron 24, 31–47 (1999).

50. Engel, A. K. & Singer, W. Temporal binding and the neural correlates of sensory awareness. Trends in Cognitive Sciences 5, 16–25 (2001).

51. Klimesch, W., Freunberger, R. & Sauseng, P. Oscillatory mechanisms of process binding in memory. Neuroscience & Biobehavioral Reviews 34, 1002–1014 (2010).

52. Senoussi, M., Verbeke, P. & Verguts, T. Time-Based Binding as a Solution to and a Limitation for Flexible Cognition. Front. Psychol. 12, 798061 (2022).

